# Linking CFTR modulators to opportunistic bacterial infections in cystic fibrosis

**DOI:** 10.1101/2022.02.15.478594

**Authors:** Cristina Cigana, Ruggero Giannella, Alice Colavolpe, Beatriz Alcalá-Franco, Giulia Mancini, Colombi Francesca, Chiara Bigogno, Ulla Bastrup, Giovanni Bertoni, Alessandra Bragonzi

## Abstract

Cystic fibrosis transmembrane conductance regulator (CFTR) modulators improve clinical outcomes with variable efficacy in patients with cystic fibrosis (CF). However, changes produced by bacterial persistence and adaptation in addition to antibiotic regimens could influence CFTR modulator efficacy and *vice versa* and hence clinical outcomes. We first evaluated the effects of ivacaftor (IVA), lumacaftor (LUM), tezacaftor, elexacaftor and elexacaftor/tezacaftor/ivacaftor (ETI), alone or combined with antibiotics, on sequential *Staphylococcus aureus* and *Pseudomonas aeruginosa* CF isolates. IVA and ETI showed the most potent direct antimicrobial activity against *S. aureus*, while *P. aeruginosa* was not affected. Additive effects or synergies were observed between the CFTR modulators and antibiotics against both *S. aureus* and *P. aeruginosa*, independently of the stage of colonization. IVA and LUM were the most effective in potentiating antibiotic activity against *S. aureus*, while IVA and ETI enhanced mainly polymyxins activity against *P. aeruginosa*. Next, we evaluated the effect of *P. aeruginosa* pneumonia on the pharmacokinetics of IVA in mice. The time-concentration curves of IVA and its metabolites in plasma, lung and epithelial lining fluid were influenced by *P. aeruginosa* infection. The area under the concentration-time curve showed that airway exposure to IVA was greater in infected than non-infected mice. These results suggest that CFTR modulators can have direct antimicrobial properties and/or enhance antibiotic activity against *S. aureus* and *P. aeruginosa*. Furthermore, bacterial infection impacts the IVA concentration and airway exposure, potentially affecting its efficacy. Our findings suggest optimizing host- and pathogen-directed drug regimens to improve efficacy for personalized treatment.

## Introduction

Cystic fibrosis (CF) is caused by mutations in the cystic fibrosis transmembrane conductance regulator (CFTR) gene and affects mostly the lung, but also other organs. Respiratory failure is caused by pulmonary infections and inflammation [1], with *Pseudomonas aeruginosa* and *Staphylococcus aureus* being recognized as the most prevalent pathogens. Their chronic persistence and pathogenicity are associated with adaptation to the CF airways [2].

Traditionally, treatments focused on lessening infection and inflammation of CF disease, although efficacy is not optimal. Recently, new approaches to correct CFTR cellular misprocessing (correctors) and restore its channel function (potentiators) have emerged [3]. The CFTR potentiator ivacaftor (IVA) was introduced to target gating and other residual function deficits [4–6], while the CFTR correctors lumacaftor (LUM), tezacaftor (TEZ) and elexacaftor (ELX) were approved to correct the F508del*-*CFTR protein and recently extended to other mutations causing processing and trafficking defects. Since the commonest F508del*-*CFTR requires both correction and potentiation for clinical efficacy, two dual- (LUM/IVA and TEZ/IVA) [7–11] and triple-(ELX/TEZ/IVA named ETI) [12] agent drugs were approved. Despite the clinical benefits of these treatments, marked variability in CF patients has been reported [4, 5, 7, 13]. This supports the view that the response to CFTR-directed therapeutics is multifactorial and that CFTR-independent factors may contribute to treatment efficacy, but the mechanisms remain unknown.

There continues to be a substantial gap in our understanding of whether and how CFTR-directed therapies impact the microbiological profile in treated patients [14, 15]. IVA treatment showed an initial reduction in the prevalence of bacterial isolation, but *P. aeruginosa* density rebounded after one year [16], and *P. aeruginosa* strains present before treatment persisted even after intensive antibiotic therapy [17]. I*n vitro* studies have shown the anti-*S. aureus* activity of IVA and its synergy with anti-*P. aeruginosa* antibiotics [18–23]. LUM and TEZ have also been shown to enhance the activity of polymyxin B (PMB) against some *P. aeruginosa* isolates [21], while the impact of ELX and ETI remains largely unexplored. Whether the activity of CFTR modulators differs in CF strains associated with early and advanced stages of lung colonization remains unclear.

To monitor possible interactions between CFTR modulators and CF pathogens, it is critical to know the drug concentration in the lung, particularly in the epithelial lining fluid (ELF), where bacteria are present [24]. Most pharmacokinetic (PK) studies have evaluated the amount of CFTR modulators and their metabolites in the plasma of different hosts [25, 26], while few data on human sputum have been reported [27]. More importantly, the amounts of CFTR modulators that reach the airway lumen and whether bacterial pneumonia can alter their PK profiles are unclear. Considering that most CF patients are likely colonized by bacteria when CFTR modulator therapy is initiated [14], there is a clinical need to assess the impact of infection on drug metabolism to determine the correct dosage. Although mouse models are not helpful for efficacy studies, they can facilitate PK analysis and provide much-needed knowledge on the interactions between CFTR modulators and CF pathogens avoiding variables of human studies.

Thus, we aimed to establish whether and how the bacterial adaptation affects the antimicrobial activity of CFTR modulators, including ELX and ETI, and the impact of bacterial infection on PK. We designed the study to i) determine the antimicrobial activity of CFTR modulators and their additive or synergistic effects with antibiotics against *P. aeruginosa* and *S. aureus* reference and clonal isolates collected from CF patients at early and advanced stages of lung colonization; ii) evaluate the concentration of IVA, as a model CFTR modulator, in plasma and airway samples and its changes during pneumonia in mice.

## Methods

### Ethics statement

The animal studies adhered to the Italian Ministry of Health guidelines for the use and care of experimental animals (IACUC #954). The experiments with CF *P. aeruginosa* and *S. aureus* isolates and storage of biological materials were approved by the Ethics Commissions of Hannover Medical School and University Hospital Münster (Germany).

### Bacterial strains

*P. aeruginosa* and *S. aureus* strains included reference PAO1 and Newman. CF clinical isolates recovered at early and late chronic colonization were RP73, AA2, AA43, AA44, MF1, MF51, KK1, KK2, KK71 and KK72 for *P. aeruginosa* and A10, A12, J6, J9, F1 and F5 [28–31] for *S. aureus*. Phenotype and antimicrobial resistance were previously characterized (**Supplementary Tables S1** and **S2**) [28–31].

### Minimum inhibitory concentration (MIC) measurement

The MICs of CFTR modulators were determined using the broth microdilution susceptibility testing method as described in Supplementary Methods [28, 32].

### Checkerboard assay

The synergistic activities of the CFTR modulators combined with tobramycin (TOB), ciprofloxacin (CIP), colistin (CST), PMB, meropenem (MER) and azithromycin (AZM) for *P. aeruginosa* and amoxicillin (AMX), teicoplanin (TEC), linezolid (LZD), vancomycin (VAN) and AZM for *S. aureus*, were determined by the checkerboard method in cation-adjusted Mueller-Hinton broth [33], as detailed in the Supplementary Methods.

### Mouse model

C57BL/6NCrlBR male mice (8-10 weeks of age) were challenged with 1×10^6^ colony forming units of the planktonic PAO1 by intratracheal injection. Mice were treated with 3 mg/kg IVA or vehicle (10% PEG 400, 10% Tween 80, and 80% saline) [19] by intraperitoneal injection half an hour after infection, according to the ARRIVE guidelines [34]. IVA dose was calculated based on a single adult dose (150 mg) adjusted for mouse weight, assuming that an adult with CF weighs 50 kg [19]. Blood, bronchoalveolar lavage fluid (BALF) and lung samples were recovered at different time points, as described [35] for PK and reported in Supplementary Methods.

### Protein binding and concentration

Protein binding was measured in mouse plasma and lung homogenate by Rapid Equilibrium Dialysis. Samples were analysed by ultra-performance liquid chromatography-tandem mass spectrometry. Quantitation was performed via multiple reaction monitoring using the transitions reported in **Supplementary Table S3**. Concentrations of IVA and its metabolites in ELF were determined by using the ratio of the urea concentration in BALF to that in plasma [36]. Protein binding is expected to be negligible in ELF [36]. The drug concentration in ELF was assumed to be equal to the unbound concentration. Additional details are reported in the Supplementary Methods.

### PK evaluation

PK parameters were estimated using PK Solver Excel using a non-compartmental approach consistent with the intraperitoneal route of IVA administration. The area under the concentration-versus-time curve was calculated using the linear trapezoidal method for the period from *t* = 0 to the time of the last quantifiable concentration level (AUC_0–last_). Evaluation of the terminal elimination phase was not practical, as terminal phase concentration data were either not available (low doses) or sparse.

### Statistics

Statistical analyses were performed with GraphPad Prism (GraphPad Software, Inc., San Diego, CA, USA) using a non-parametric two-tailed Mann-Whitney U test. A value of p ≤ 0.05 was considered statistically significant.

## Results

### IVA and ETI show the most potent antimicrobial activity against longitudinal *S. aureus* isolates, while *P. aeruginosa* isolates are not affected

To evaluate the impact of the bacterial adaptation on the antimicrobial activity of CFTR modulators we measured MICs against clinical clonal isolates, collected from CF patients at early and late stage of chronic colonization (**Supplementary Tables S1** and **S2**) [28–31]. Single CFTR modulators and the ELX/TEZ/IVA combination at a ratio of 2/1/1.5, which is used for CF patients [37], were tested. IVA showed antimicrobial activity with MICs ranging from 2 to 8 μg/ml for all the *S. aureus* isolates (**Table 1**). The MIC of ELX was 16 μg/ml for almost all *S. aureus* isolates, except for the J6 isolate (32 μg/ml). By contrast, the MICs of LUM and TEZ were >32 μg/ml, indicating little or no effect against any of the *S. aureus* isolates. The ELX/TEZ/IVA combination showed antimicrobial activity against all the *S. aureus* isolates, with MICs ranging from 4/2/3 to 8/4/6 μg/ml. By contrast, all CFTR modulators showed MICs >32 μg/ml against *P. aeruginosa* isolates (**Supplementary Table S4**). Our results demonstrate that adaptation does not change either the susceptibility of *S. aureus* strains to IVA and ETI or the resistance of *P. aeruginosa* to CFTR modulators.

**Table 1.**
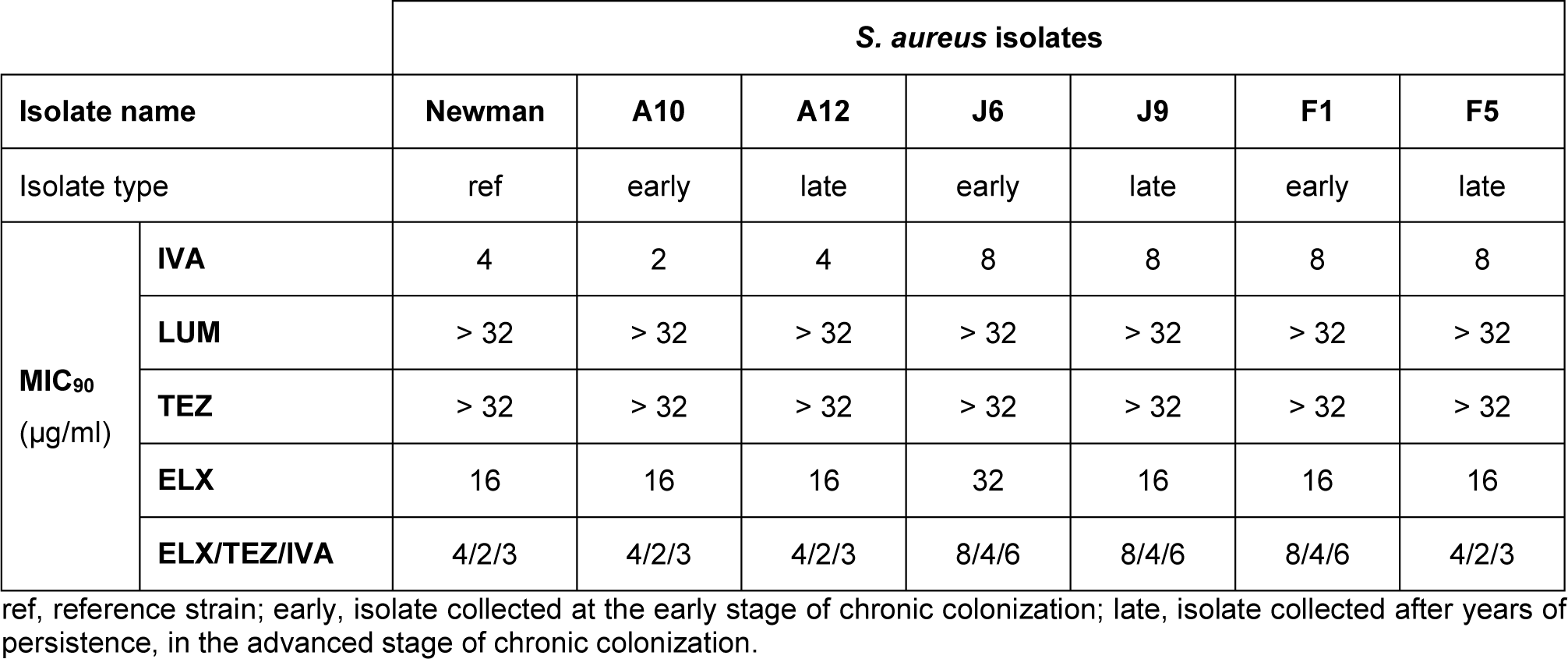
Minimum inhibitory concentrations (MIC_90_) of ivacaftor (IVA), lumacaftor (LUM), tezacaftor (TEZ), elexacaftor (ELX) and the triple combination elexacaftor/tezacaftor/ivacaftor (ELX/TEZ/IVA) against *S. aureus* isolates collected from CF patients at different stages of colonization (early and late). *S. aureus* isolates were grown for 20 hrs in the presence of serial dilutions of IVA, LUM, TEZ, ELX and the triple combination ELX/TEZ/IVA. The concentrations tested ranged from 0.25 μg/ml to 32 μg/ml for IVA, LUM, TEZ and ELX and from ELX 0.5 μg/ml/TEZ 0.25 μg/ml/IVA 0.375 μg/ml to ELX 64 μg/ml/TEZ 32 μg/ml/IVA 48 μg/ml for the triple combination. The MIC_90_ was defined as the lowest compound concentration showing a reduction in the optical density at 620 nm of approximately 90% in comparison to the optical density of the bacteria grown with the vehicle after 20 hrs. Each experiment was performed at least two independent times (two technical replicates).

|  |  | *S. aureus* isolates |  |  |  |  |  |  |
| --- | --- | --- | --- | --- | --- | --- | --- | --- |
| Isolate name |  | Newman | A10 | A12 | J6 | J9 | F1 | F5 |
| Isolate type |  | ref | early | late | early | late | early | late |
| MIC<sub>90</sub><br>(µg/ml) | IVA | 4 | 2 | 4 | 8 | 8 | 8 | 8 |
|  | LUM | > 32 | > 32 | > 32 | > 32 | > 32 | > 32 | > 32 |
|  | TEZ | > 32 | > 32 | > 32 | > 32 | > 32 | > 32 | > 32 |
|  | ELX | 16 | 16 | 16 | 32 | 16 | 16 | 16 |
|  | ELX/TEZ/IVA | 4/2/3 | 4/2/3 | 4/2/3 | 8/4/6 | 8/4/6 | 8/4/6 | 4/2/3 |

ref, reference strain; early, isolate collected at the early stage of chronic colonization; late, isolate collected after years of persistence, in the advanced stage of chronic colonization.

### IVA and LUM potentiate the activity of all antibiotics against *S. aureus*

Next, we determined potential additive or synergistic effects of the CFTR modulators in combination with antibiotics against reference and clinical *S. aureus* isolates by checkerboard assays. In general, except for TEZ, interaction effects were observed on some isolates and regardless of the stage of colonization (**Table 2**). The fractional inhibitory concentration (FIC) indexes of IVA showed broad-spectrum potentiation of LZD activity (6 out of 7 isolates), while a narrower spectrum was observed for AMX, VAN, and TEC. Synergistic effects were observed also between IVA and either VAN or TEC but only in the Newman reference strain. By contrast, additive effects between either ELX or ETI and the tested antibiotics were more sporadic or completely absent, as in the case of LZD. LUM, which did not show direct antibiotic effects, behaved similarly to IVA in elevating the activities of the tested antibiotics and showed better synergistic effects with VAN or TEC against the Newman strain. These findings indicate that the CFTR modulators, mainly IVA and LUM, can enhance the performance of antibiotics used against *S. aureus* infections in CF patients. The potentiating effects appear to be bacterial isolate dependent, but no correlation with the stage of chronic colonization was found.

**Table 2.** Fractional inhibitory concentration (FIC) indexes for ivacaftor (IVA), lumacaftor (LUM), tezacaftor (TEZ), elexacaftor (ELX) and the triple combination elexacaftor/tezacaftor/ivacaftor (ELX/TEZ/IVA) in combination with common antibiotics against *S. aureus* isolates collected from CF patients. *S. aureus* isolates were grown for 20 hrs in the presence of serial dilutions of IVA, LUM, TEZ, ELX or the triple combination ELX/TEZ/IVA (ETI) and amoxicillin (AMX), vancomycin (VAN), teicoplanin (TEC), linezolid (LZD), or azithromycin (AZM). The concentrations of the CFTR modulators tested ranged from 0.5 μg/ml to 32 μg/ml for IVA, LUM and ELX and from ELX 0.5 μg/ml/TEZ 0.25 μg/ml/IVA 0.375 μg/ml to ELX 32 μg/ml/TEZ 16 μg/ml/IVA 24 μg/ml for the triple combination ETI, while those of the antibiotics started from the concentration two-fold greater than the MIC_90_ for the isolate to 32 times less than the MIC_90_ of the antibiotic alone. FIC indexes, calculated as MIC_90_ of CFTR modulator in combination/ MIC_90_ CFTR alone + MIC_90_ antibiotic in combination/ MIC90 antibiotic alone, are indicated. Each experiment was performed at least two independent times (two technical replicates).

|  |  |  | *S. aureus* isolates |  |  |  |  |  |  |
| --- | --- | --- | --- | --- | --- | --- | --- | --- | --- |
|  |  |  | Newman | A10 | A12 | J6 | J9 | F1 | F5 |
|  |  |  | ref | early | late | early | late | early | late |
| FIC<br>index* | IVA | AMX | 0.5 | 1 | 2 | 1 | 0.5 | 0.625 | 0.625 |
|  |  | VAN | 0.31 | 0.625 | 0.625 | 1 | 1 | 1 | 1 |
|  |  | TEC | 0.31 | 1 | 1 | 1 | 0.75 | 1 | 0.75 |
|  |  | LZD | 0.75 | 2 | 0.625 | 0.625 | 0.625 | 0.625 | 0.625 |
|  |  | AZM | 0.531 | 2 | 2 | 2 | 2 | 2 | 2 |
|  | LUM | AMX | 0.625 | 0.625 | 2 | 1 | 0.625 | 0.625 | 0.625 |
|  |  | VAN | 0.155 | 0.625 | 0.75 | 1 | 1 | 1 | 0.625 |
|  |  | TEC | 0.185 | 0.625 | 0.625 | 2 | 2 | 0.625 | 0.75 |
|  |  | LZD | 0.625 | 2 | 0.625 | 2 | 0.75 | 0.625 | 0.625 |
|  |  | AZM | 0.75 | 1 | 1 | 2 | 2 | 1 | 2 |
|  | TEZ | AMX | 2 | 2 | 2 | 2 | 2 | 2 | 2 |
|  |  | VAN | 2 | 2 | 2 | 2 | 2 | 2 | 2 |
|  |  | TEC | 2 | 2 | 2 | 2 | 2 | 2 | 2 |
|  |  | LZD | 2 | 2 | 2 | 2 | 2 | 2 | 2 |
|  |  | AZM | 2 | 2 | 2 | 2 | 1 | 2 | 2 |
|  | ELX | AMX | 2 | 0.5 | 0.75 | 2 | 0.75 | 2 | 2 |
|  |  | VAN | 2 | 0.563 | 2 | 2 | 2 | 2 | 2 |
|  |  | TEC | 2 | 2 | 2 | 2 | 2 | 2 | 2 |
|  |  | LZD | 2 | 2 | 2 | 2 | 2 | 2 | 2 |
|  |  | AZM | 0.75 | 1 | 1 | 0.75 | 0.75 | 0.5 | 2 |
|  | ETI | AMX | 2 | 0.75 | 1 | 2 | 0.75 | 2 | 2 |
|  |  | VAN | 2 | 2 | 2 | 2 | 2 | 2 | 2 |
|  |  | TEC | 2 | 2 | 2 | 0.625 | 2 | 1 | 2 |
|  |  | LZD | 2 | 2 | 2 | 2 | 2 | 2 | 2 |
|  |  | AZM | 2 | 2 | 2 | 2 | 2 | 2 | 2 |

\*FIC index <0.5 indicates synergy (highlighted in green), FIC index >0.50 and <1.0 indicates additive effect (highlighted in yellow), FIC index >1.0 and <4 indicates indifference. Ref, reference strain; early, isolate collected at the early stage of chronic colonization; late, isolate collected after years of persistence, in the advanced stage of chronic colonization.

### IVA and ETI potentiate the activity of only selected antibiotics against *P. aeruginosa*

Despite the lack of intrinsic antibacterial activity of the CFTR modulators against *P. aeruginosa*, we tested whether they enhance the activity of antibiotics (**Table 3**). Strikingly, IVA was found to strongly enhance the activity of CST and PMB against 95.45% of the isolates. Synergy, rather than additivity, was detected in most of the isolates. No effect was observed when IVA was combined with CIP, MER or TOB. LUM, TEZ and ELX showed only a few cases of additive or synergistic effects with selected antibiotics. For ETI, the FIC values indicated an additive effect on the activities of CST and PMB in several isolates, with synergy against the AA2 isolate reached with both antibiotics. In only two isolates, ETI potentiated the activity of CIP. No CFTR modulators affected AZM activity on *P. aeruginosa* isolates. Notably, no negative interaction of the CFTR modulators with the antibiotics was observed. These findings indicate that IVA and ETI can enhance antibiotic activity, particularly that of CST and PMB, against *P. aeruginosa* in an isolate-dependent manner and independently of the stage of colonization.

**Table 3.** Fractional inhibitory concentration (FIC) indexes for ivacaftor (IVA), lumacaftor (LUM), tezacaftor (TEZ), elexacaftor (ELX) and the triple combination elexacaftor/tezacaftor/ ivacaftor (ELX/TEZ/IVA) in combination with common antibiotics against *P. aeruginosa* isolates collected from CF patients. *P. aeruginosa* isolates were grown for 20 hrs in the presence of serial dilutions of IVA, LUM, TEZ, ELX or the triple combination ELX/TEZ/IVA and ciprofloxacin (CIP), meropenem (MER), colistin (CST), tobramycin (TOB), polymyxin B (PMB) and azithromycin (AZM). The concentrations of the CFTR modulators tested ranged from 0.5 μg/ml to 32 μg/ml for IVA, LUM, TEZ and ELX, and from ELX 0.5 μg/ml/TEZ 0.25 μg/ml/IVA 0.375 μg/ml to ELX 32 μg/ ml/TEZ 16 μg/ml/IVA 24 μg/ml for the triple combination ETI, while those of the antibiotics started from the concentration two-fold greater than the MIC_90_ for the isolate to 32 times less than the MIC_90_ of the antibiotic alone. FIC indexes, calculated as MIC_90_ of CFTR modulator in combination/ MIC_90_ CFTR alone + MIC_90_ antibiotic in combination/ MIC_90_ antibiotic alone, are indicated. Each experiment was performed at least two independent times (two technical replicates).

|  |  |  | *P. aeruginosa* isolates |  |  |  |  |  |  |  |  |  |  |
| --- | --- | --- | --- | --- | --- | --- | --- | --- | --- | --- | --- | --- | --- |
|  |  |  | PAO1 | AA2 | AA43 | AA44 | MF1 | MF51 | KK1 | KK2 | KK71 | KK72 | RP73 |
|  |  |  | ref | early | late | late | early | late | early | early | late | late | late |
| FIC index* | IVA | CIP | 2 | 2 | 2 | 2 | 2 | 2 | 2 | 2 | 2 | 2 | 2 |
|  |  | MER | 2 | 2 | 2 | 2 | 2 | 2 | 2 | 2 | 2 | 2 | 2 |
|  |  | TOB | 2 | 2 | 2 | 2 | 2 | 2 | 2 | 2 | 2 | 2 | 2 |
|  |  | CST | 0.531 | 0.156 | 0.281 | 0.281 | 0.531 | 0.531 | 2 | 0.531 | 0.312 | 0.281 | 0.281 |
|  |  | PMB | 0.562 | 0.281 | 0.281 | 0.281 | 0.281 | 0.281 | 0.375 | 0.516 | 0.312 | 0.281 | 0.281 |
|  |  | AZM | 2 | 2 | 2 | 2 | 2 | 2 | 2 | 2 | 2 | 2 | 2 |
|  | LUM | CIP | 2 | 0.625 | 2 | 0.5 | 0.281 | 0.75 | 0.625 | 1 | 2 | 2 | 2 |
|  |  | MER | 2 | 2 | 2 | 0.5 | 2 | 2 | 2 | 1 | 2 | 2 | 2 |
|  |  | TOB | 1 | 2 | 2 | 0.5 | 2 | 2 | 1 | 0.75 | 1 | 1 | 0.75 |
|  |  | CST | 1 | 0.625 | 2 | 2 | 2 | 2 | 2 | 2 | 0.5 | 0.375 | 2 |
|  |  | PMB | 2 | 2 | 2 | 2 | 2 | 2 | 2 | 2 | 2 | 2 | 2 |
|  |  | AZM | 2 | 2 | 2 | 2 | 2 | 2 | 2 | 2 | 2 | 2 | 2 |
|  | TEZ | CIP | 2 | 2 | 2 | 2 | 2 | 2 | 2 | 2 | 2 | 2 | 2 |
|  |  | MER | 2 | 2 | 2 | 2 | 2 | 2 | 2 | 2 | 2 | 2 | 2 |
|  |  | TOB | 2 | 2 | 2 | 2 | 2 | 2 | 2 | 2 | 2 | 2 | 2 |
|  |  | CST | 2 | 2 | 2 | 2 | 2 | 2 | 2 | 0.75 | 2 | 2 | 2 |
|  |  | PMB | 2 | 0.562 | 2 | 2 | 2 | 2 | 2 | 2 | 2 | 2 | 2 |
|  |  | AZM | 2 | 2 | 2 | 2 | 2 | 2 | 2 | 2 | 2 | 2 | 2 |
|  | ELX | CIP | 2 | 2 | 2 | 2 | 0.687 | 2 | 2 | 2 | 2 | 2 | 2 |
|  |  | MER | 2 | 2 | 2 | 2 | 2 | 1 | 2 | 2 | 2 | 2 | 2 |
|  |  | TOB | 2 | 2 | 2 | 2 | 2 | 2 | 2 | 2 | 2 | 2 | 2 |
|  |  | CST | 2 | 0.562 | 2 | 2 | 2 | 2 | 2 | 2 | 2 | 2 | 2 |
|  |  | PMB | 2 | 2 | 2 | 2 | 2 | 2 | 2 | 2 | 2 | 2 | 2 |
|  |  | AZM | 2 | 2 | 2 | 2 | 2 | 2 | 2 | 2 | 2 | 2 | 2 |
|  | ETI | CIP | 2 | 2 | 2 | 2 | 0.438 | 1.25 | 2 | 0.594 | 1.25 | 2 | 2 |
|  |  | MER | 2 | 2 | 2 | 2 | 2 | 2 | 2 | 2 | 2 | 2 | 2 |
|  |  | TOB | 2 | 2 | 2 | 2 | 2 | 2 | 2 | 2 | 2 | 2 | 2 |
|  |  | CST | 0.523 | 0.273 | 0.523 | 0.547 | 0.547 | 2 | 2 | 2 | 0.594 | 0.547 | 2 |
|  |  | PMB | 2 | 0.297 | 0.512 | 2 | 2 | 0.547 | 2 | 2 | 0.547 | 0.547 | 2 |
|  |  | AZM | 2 | 2 | 2 | 2 | 2 | 2 | 2 | 2 | 2 | 2 | 2 |

\*FIC index <0.5 indicates synergy (highlighted in green), FIC index >0.50 and <1.0 indicates additive effect (highlighted in yellow), FIC index >1.0 and <4 indicates indifference. Ref, reference strain; early, isolate collected at the early stage of chronic colonization; late, isolate collected after years of persistence, in the advanced stage of chronic colonization.

### IVA is differentially distributed in plasma, lung and ELF, with concentrations affected by *P. aeruginosa* infection

To establish the impact of bacterial infection on CFTR modulators PK, we exploited the mouse model of *P. aeruginosa* pneumonia [35] and evaluated IVA concentration in murine plasma, lung and ELF. We selected IVA as the CFTR modulator with the most striking anti-microbial activity and *P. aeruginosa* as the most relevant CF pathogen [38]. We first determined the protein binding of IVA and its metabolites M1 (pharmacologically active) and M6 (inactive) [39]. The percentages of protein binding were very high in the plasma, with those of IVA, M1 and M6 being ≥99.5% (**Table 4**). Similar protein binding was observed in the lung for IVA (99.7%) and M1 (97.2%) while that of M6 was lower (80.3%).

**Table 4.** Percentages of ivacaftor (IVA) and its metabolites (M1 and M6) as protein-bound and free active compounds in murine plasma and lungs. Blood was collected from C57BL/6NCrlBR male mice (8 to 10 weeks of age) and processed to obtain plasma. Lungs were excised, homogenized, and centrifuged, and the supernatants were collected. Protein binding was measured by the dialysis method with Rapid Equilibrium Dialysis inserts, with undiluted plasma or lung homogenate from 2-hr samples derived from non-infected untreated mice diluted 1:2 in PBS buffer spiked with a solution containing the three analytes against PBS buffer. Samples were analysed with ultra-performance liquid chromatography-tandem mass spectrometry in positive multiple reaction monitoring mode. Values are the means of three replicates.

| Sample |  | Protein binding (%) |  |  |
| --- | --- | --- | --- | --- |
|  |  | IVA | M1 | M6 |
| Plasma | Bound fraction | 99.97 | 99.89 | 99.47 |
|  | Free active fraction | 0.03 | 0.11 | 0.53 |
| Lung | Bound fraction | 99.74 | 97.17 | 80.3 |
|  | Free active fraction | 0.26 | 2.83 | 19.7 |

Next, the time-concentration curves of IVA and its metabolites showed that parent drug accounted for the majority of the total drug, followed by M1 and then M6, in the plasma of both infected and non-infected mice (**Figure 1A**). At early time points, IVA and M1 levels were lower in *P. aeruginosa*-infected compared to non-infected mice (p=0.057). However, at later time points, the IVA levels were slightly higher in infected compared to non-infected mice. M6 levels ranged from low to undetectable in the plasma of all mice. Peak plasma levels of IVA, reached 1 hr after treatment, were higher in non-infected compared to infected mice (**Table 5**), as were M1 levels (**Supplementary Table S5**). Nonetheless, the AUC for the period from t_10min_ to the last quantifiable concentration (AUC_last_) and the AUC to infinity (AUC_inf_) for IVA were higher in *P. aeruginosa*-infected than non-infected mice, indicating higher exposure in the presence of infection. This was confirmed by higher last quantifiable concentration (C_last_) at 24 hrs, half-life (T_1/2_) and mean residence time (MRT) in infected compared to non-infected mice. By contrast, M1 AUCs, C_last_, T_1/2_ and MRT were similar in both groups.

**Table 5.** Pharmacokinetic parameter estimates for ivacaftor (IVA) in plasma, epithelial lining fluid (ELF) and lungs from infected and non-infected mice. C57BL/6NCrlBR male mice (8 to 10 weeks of age) were infected with 1×10^6^ colony forming units of *P. aeruginosa* PAO1 by intratracheal administration. A non-infected control group was also tested in parallel. Thirty minutes after infection, the mice were treated with 3 mg/kg IVA in 10% PEG 400, 10% Tween 80, and 80% saline by intraperitoneal administration. Mice were sacrificed at 10 min and 1, 2, 6 and 24 hrs after IVA administration. Blood was collected and processed to obtain plasma. Bronchoalveolar lavage fluid (BALF) was collected and centrifuged, and the supernatant was used to quantify the IVA concentration. Lungs were excised, homogenized, and centrifuged, and the supernatants were used to quantify IVA concentrations. Plasma, lung homogenate and BALF were added to a Phree phospholipid removal plate (Phenomenex) with acetonitrile and 0.1% formic acid in order to eliminate phospholipids, decreasing the matrix effect. Eluates were analysed by the ultra-performance liquid chromatography-tandem mass spectrometry method with a linear gradient in multiple reaction monitoring positive mode. Calibration ranges were 0.5-750 ng/ml in plasma and BALF and 4-1000 ng/g in the lung homogenate. Concentrations of IVA in ELF were determined using the ratio of the urea concentration in plasma to that in BALF. Concentration in ELF = drug concentration in BALF × urea in plasma / urea in BALF. The data are the geometric means of values from 3-4 mice.

| Sample | Mouse group | Pharmacokinetic parameters |  |  |  |  |  |  |  |
| --- | --- | --- | --- | --- | --- | --- | --- | --- | --- |
|  |  | T<sub>max</sub> (hrs) | C<sub>max</sub> (ng/ml) | T<sub>last</sub> (hrs) | C<sub>last</sub> (hrs) | AUC<sub>last</sub> (hrs ng/ml) | AUC<sub>IN</sub> F (hrs ng/ml) | T<sub>1/2</sub> (hrs) | MRT (hrs) |
| Plasma | non-infected | 1 | 2345 | 24 | 4.0 | 7482 | 7498 | 2.9 | 2.9 |
|  | infected | 1 | 968 | 24 | 8.6 | 9111 | 9152 | 3.3 | 4.7 |
| ELF | non-infected | 1 | 537 | 6 | 116.7 | 1932 | 2307 | 2.2 | 2.1 |
|  | infected | 1 | 981 | 6 | 194.3 | 2590 | 3236 | 2.3 | 2.2 |
| Lung | non-infected | 1 | 249 | 6 | 68.2 | 946 | nd | nd | 2.2 |
|  | infected | 1 | 364 | 24 | 4.4 | 4010 | 4033 | 3.7 | 5.2 |

T<sub>max</sub>, time of maximum concentration; C<sub>max</sub>, maximum concentration; T<sub>last</sub>, time of last quantifiable concentration; C<sub>last</sub>, last quantifiable concentration; AUC<sub>last</sub>, AUC to the last quantifiable concentration level; AUC<sub>INF</sub>, AUC to infinity; T<sub>1/2</sub>, half-life; MRT, mean residence time; nd, not determined.

**Figure 1.**
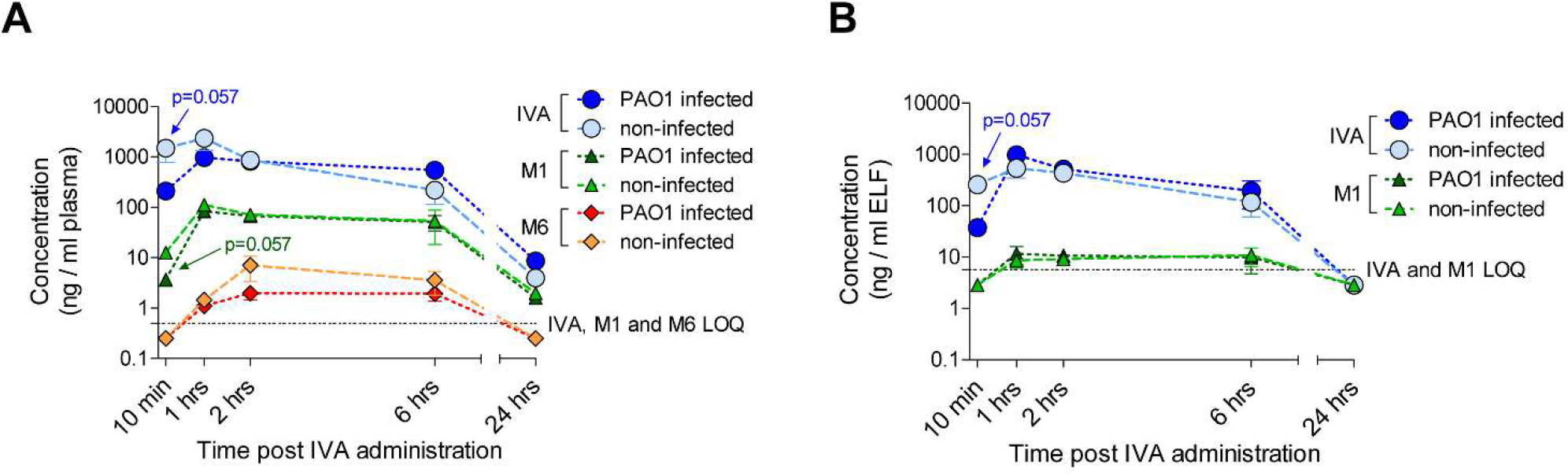
Ivacaftor (IVA), M1 and M6 concentrations in murine plasma and epithelial lining fluid (ELF). C57BL/6NCrlBR male mice (8 to 10 weeks of age) were infected with 1×10^6^ colony forming units of *P. aeruginosa* PAO1 by intratracheal administration. A non-infected control group was also tested in parallel. Thirty minutes after infection, the mice were treated with 3 mg/kg IVA in 10% PEG 400, 10% Tween 80, and 80% saline by intraperitoneal administration. Mice were sacrificed at 10 min and 1, 2, 6 and 24 hrs after IVA administration. Blood was collected and processed to obtain plasma. IVA, M1 and M6 concentrations in plasma were measured by high-performance liquid chromatography–tandem mass spectrometry (**A**). Bronchoalveolar lavage fluid (BALF) was collected and centrifuged, and the supernatant was used to quantify the IVA concentration. The volume of the ELF was determined by using the ratio of the urea concentration in the BALF to that in the plasma (**B**). The amounts of IVA and M1 were measured by high-performance liquid chromatography–tandem mass spectrometry. Data (derived from 3-4 mice) are represented as the mean values ± standard errors of the means (SEMs). Limits of quantification (LOQ) for IVA and its metabolites are indicated. A value corresponding to LOQ/2 was assigned to undetectable samples at specific time-points (e.g. 10 min and/or 24 hrs). Statistical significance was calculated by the Mann-Whitney test comparing the infected and non-infected mouse groups at each time point.

The IVA profile in the ELF followed that in the plasma samples up to 6 hrs, with higher levels in non-infected compared to infected mice soon after treatment (p=0.057; **Figure 1B**). *P. aeruginosa* infection resulted in higher IVA peak levels, AUC_last_, AUC_inf_ and C_last_ compared to non-infected mice (**Table 5**). The partitioning to the ELF was estimated by comparing the AUC of the unbound compound (fAUC) in the plasma to that in the ELF. For the calculation of fAUC, a factor of 0.03% was used for the IVA parent drug in the plasma (**Table 4**). The ELF/plasma fAUC ratio of IVA was high, indicating that IVA penetrates ELF well (**Table 6**). M1 was only detected between 1 and 6 hrs post-administration in ELF of infected and non-infected mice (**Figure 1B**). The ELF/plasma fAUC ratio was also high for M1, indicating high penetration in the ELF. M6 was not quantifiable for any conditions.

**Table 6.** Penetration of ivacaftor (IVA) in epithelial lining fluid (ELF) and lungs. The C_max_ and AUC of plasma and lung homogenates were corrected for the free fraction according to the protein binding experiment (0.03% for plasma and 0.26% for lung; **Table 4**) to obtain the C_max_ and AUC of unbound IVA (fC_max_ and fAUC). Data are the geometric means of values from 3-4 mice.

| Mouse group | Values by sample type |  |  |  |  |  | Lung/plasma |  | ELF/plasma |  |
| --- | --- | --- | --- | --- | --- | --- | --- | --- | --- | --- |
|  | plasma |  | lung |  | ELF |  |  |  |  |  |
|  | fC<sub>max</sub><br>(ng/ml<br>) | fAUC<br>(hrs<br>ng/ml) | fC<sub>max</sub><br>(ng/ml<br>) | fAUC<br>(hrs<br>ng/ml) | C<sub>max</sub><br>(ng/ml<br>) | AUC<br>(hrs<br>ng/ml) | C<sub>max</sub><sup>#</sup> | AUC<sup>#</sup> | C<sub>max</sub><sup>#</sup> | AUC<sup>#</sup> |
| Non-infected | 0.70 | 2.20 | 0.65 | 2.44 | 537 | 1910 | 0.93 | 1.11 | 767 | 868 |
| Infected | 0.29 | 2.72 | 0.94 | 10.62 | 981 | 2587 | 3.24 | 3.90 | 3383 | 951 |

<sup>#</sup> Values for lung/plasma (ratio of lung free maximum concentration or lung exposure to free maximum concentration divided by those in plasma) and ELF/plasma (ratio of ELF maximum concentration or ELF exposure to maximum concentration divided by plasma free maximum concentration or plasma exposure to free maximum concentration); fC<sub>max</sub> and fAUC refer to unbound IVA in the plasma and lung.

The IVA profile in the lung homogenates showed similar levels in infected and non-infected mice up to 2 hrs, while the levels were higher in *P. aeruginosa*-infected mice after 6 hrs (**Supplementary Figure S1**). The lung/plasma fAUC ratio of IVA indicated that IVA penetrates the lung very well, particularly in infected mice (**Table 6**). M1 showed a profile in the lung similar to that in the ELF in both infected and non-infected mice (**Supplementary Figure S1** and **Supplementary Tables S5 and S6**). M6 was not quantifiable in either infected or non-infected mice at any time point.

These findings indicated that the IVA parent drug, accounting for the majority of the total drug, readily distributes to the airways and is higher during *P. aeruginosa* infection.

## Discussion

CFTR modulators correct the molecular defect underlying CF and disease manifestations. Since bacterial lung infections/colonization are one of the hallmarks of CF disease, the effect (if any) of CFTR modulators on bacteria could deeply affect the course of the disease. Here, we tested the impact of bacterial adaptation on the antimicrobial activity of CFTR modulators and that of bacterial infection on PK. We found that: 1) IVA and newly licensed ETI have the most potent intrinsic antimicrobial activity against longitudinal *S. aureus* isolates, while *P. aeruginosa* isolates are not affected; 2) IVA and LUM potentiate the activity of all antibiotics against *S. aureus,* while IVA and ETI especially that of polymyxins against *P. aeruginosa;* 3) antimicrobial activity of CFTR modulators is independent of the stage of colonization for both bacterial species; 4) *P. aeruginosa* infection affects the distribution of IVA to plasma, lung and ELF.

Our study showed that IVA has a spectrum of relevant antibacterial activity that spans both early and late isolates of *S. aureus*. In addition, ETI is active against *S. aureus* and its activity seems to be mainly driven by IVA since the effective concentration of IVA in the triple combination was in the range of that as a single drug. In addition, we showed for the first time that ELX has broad antibacterial activity against *S. aureus*. Notably, the anti-*S. aureus* activity of CFTR modulators is independent of the stage of colonization. These activities deserve further investigation with additional biobanks, including those from CF patients under CFTR modulators treatment. Regarding CFTR modulator and antibiotic interactions, we expanded the notion that IVA can increase the activity of several classes of antibiotics against *S. aureus*. Furthermore, we show that LUM, with no intrinsic antibacterial activity against *S. aureus*, can positively interact with several classes of antibiotics. The specific mechanism underlying the enhancement of antibiotics activity by IVA and LUM against *S. aureus* is unknown and should be investigated.

We observed a different scenario for *P. aeruginosa*. IVA showed no antibacterial activity against any *P. aeruginosa* early and late isolates, suggesting that adaptation to the CF environment does not induce modification of bacterial structures or functions potentially causing resistance to IVA. This is in line with other reports showing that IVA is inactive against *P. aeruginosa* and other gram-negative species (e.g. *Klebsiella pneumoniae* and *Acinetobacter baumannii*) [21]. Gram-negative bacteria have a formidable outer membrane barrier, particularly against hydrophobic drugs as CFTR modulators, that can explain the absence of IVA activity. Alternatively, IVA might not affect essential *P. aeruginosa* pathways and can be effective only as a combined treatment. Indeed, we observed that IVA can specifically act in concert with polymyxins, such as CST and PMB, to enhance their antibacterial activity in almost all the isolates tested, while no additive or synergistic effects were observed for the other classes of antibiotics. Recently, IVA in combination with PMB has been shown to potentiate bactericidal activity against *P. aeruginosa* by inhibiting cell envelope biogenesis [23]. Similar to IVA, also ETI potentiated the activities of CST and PMB in several isolates. Although the activity of ETI can be driven by IVA, some exceptions have been observed, suggesting potential antagonisms when IVA is combined in the triple combination in an isolate-dependent manner.

To further interpret these results, it is critical to know the amounts of CFTR modulators that reach the airway lumen and whether bacterial pneumonia can alter their concentrations. We focused on IVA since this CFTR modulator exhibited the most effective antimicrobial activity independently or in combination with antibiotics. To investigate the PK profile and provide insight on the interactions between CFTR modulators and CF pathogens in the absence of confounding variables of human studies, we exploited a mouse model of *P. aeruginosa* pneumonia. Our results showed that the parent drug accounted for the majority of the total drug in plasma, lung and ELF, similar to previous reports focused solely on murine plasma but different from humans, who show higher concentrations of M1 in plasma [25]. Interestingly, exposure of both plasma and airways to IVA was higher in mice with acute *P. aeruginosa* lung infection than in non-infected mice. In our study, the IVA C_max_ in the plasma was in the range of a few μg/ml. However, the protein binding of IVA and its metabolites was extremely high in both the plasma and lung, indicating that the amount of free active compound is low, in agreement with previous studies [25]. For instance, the C_max_ of free active IVA in plasma was in the range of approximately 1 ng/ml in our model, similarly to the peak free plasma concentration measured in CF patients treated with IVA [26].

When we focused on the airways, IVA showed good penetration into the ELF, reaching higher concentrations than those in the plasma. The IVA C_max_ was approximately 0.5-1 μg/ml in the ELF. Notably, we observed antimicrobial activity or additive/synergistic effects with CST at IVA concentrations ranging from 1 to 8 μg/ml, which were comparable to or higher than those observed in the murine ELF. Schneider and colleagues measured an IVA concentration of 0.15±0.05 μg/ml in the sputum of a CF patient at 2.5 hrs post IVA administration [27], which would exclude antimicrobial activities of CFTR modulators in CF patients. However, data from sputa of single patients cannot be considered conclusive, and IVA quantification in a cohort of patients is needed. In addition, IVA is known to accumulate with repeated doses, particularly when it is taken with a fatty meal [40]. Our work analysed IVA concentrations in mice after administering a single dose, but higher levels are expected during chronic treatment. In this regard, IVA levels following prolonged treatment in CF patients are unknown. Therefore, the antimicrobial activities of IVA in the airways of CF patients could be plausible and deserves further investigation. In addition our findings support performing new studies on the potential benefits of pulmonary administration of CST in combination with IVA in treating *P. aeruginosa* lung infections [41].

## Conclusions

This work supports the interaction of CFTR-modulators with opportunistic bacterial infection and antibiotics. Our results underline that CFTR-modulators influence antibiotic efficacy through a direct or synergistic effect. This may orient the selection of specific antibiotics to treat infection in combination with CFTR-modulators. Importantly, CFTR-modulator efficacy is not influenced by specific bacterial phenotypes (early vs late adapted), indicating drug targeting at any stage of colonization. Since the bacterial strains used in this study were collected several years ago, they do not show some of the antibiotic resistances that have emerged in CF clinics in recent years. The challenge ahead is to validate our results with additional biobanks that include longitudinal strains isolated from CF patients under CFTR modulator therapies and with different patterns of antibiotic resistance.

So far, both animal and human studies did not clarify the impact of infection on concentrations and biodistribution of CFTR modulators. Our results in a mouse model of *P. aeruginosa* pneumonia underline the importance of testing PK during infection, particularly in the airways. This would help to optimize the drug regimens improving the efficacy in the view of personalized treatment.

## Acknowledgements

We thank Marzia Giustra for her invaluable contribution to the *in vitro* assays, and Burkhard Tümmler (Medizinische Hochschule Hannover, Germany) and Barbara C. Kahl (University Hospital Münster, Germany) for supplying respectively *P. aeruginosa* and *S. aureus* clinical isolates collected from CF patients. This study was supported to CC by the Italian Cystic Fibrosis Research Foundation (FFC#15/2018), with the contribution of the Delegazione FFC di Milano and Gruppo di Sostegno FFC di Morbegno, and to AB by the Cystic Fibrosis Foundation (BRAGON19G0). The funders had no role in study design, data collection and analysis, decision to publish, or preparation of the manuscript.

## Author contribution

CC and BA contributed to the conception and design of the study. GR, CA and MG performed MIC and checkerboard assays. CC, AFB and CF performed *in vivo* experiments and collected data. BC and BU evaluated protein binding, and compound quantification in murine samples. CC analyzed data and performed the statistical analysis. CC and BC prepared the figures and tables. CC wrote the first draft of the manuscript. BG and BA critically revised the manuscript. All the authors read, revised and approved the final manuscript.

