## Supplementary material for "Linking CFTR modulators to opportunistic bacterial infections in cystic fibrosis": CFTR modulators_Supplementary information.pdf

#### Supplementary Methods

**Minimum inhibitory concentration (MIC) measurement.** An aliquot of *P. aeruginosa* or *S. aureus* strains from glycerol stocks was streaked for isolation on tryptic soy agar (TSA) and incubated at 37°C overnight (O/N). Bacterial glycerol stocks were not used more than three times to avoid variability in the experiments. One colony was picked from the plate and used to inoculate 5 ml of tryptic soy broth (TSB) (BD, Becton and Dickinson) and placed in a shaking incubator at 37°C 200 rpm O/N. The O/N bacterial suspension was diluted to 0.1 OD/ml in 20 ml of TSB / flask and grown at 37°C at 200rpm up to reach the log phase [1]. The bacteria were pelleted by centrifugation (2700 g, 15 min, 4°C) and resuspended in cation-adjusted Mueller-Hinton broth (MH-II broth). MICs of CFTR modulators were determined using the broth microdilution susceptibility testing method, following the Clinical and Laboratory Standards Institute guidelines [2, 3]. CFTR modulators were prepared using stock solutions in dimethyl sulfoxide (DMSO), and then tested in two-fold serial diluted fashion in MH-II broth. MIC testing was run in sterile 96-well Microtiter plates. The MIC was defined as the lowest compound concentration showing a reduction in the optical density at 620 nm of approximately 90% ($MIC_{90}$) in comparison to the optical density of the bacteria grown with the vehicle after 20 hrs.

**Checkerboard assay.** The synergistic activities of CFTR modulators combined with antibiotics were determined by the checkerboard method in MH-II broth [4]. Stock solutions of CFTR modulators were prepared in DMSO, while those of antibiotics in sterile $H_2O$ or DMSO, according to their solubility. Serial dilutions of CFTR modulators and antibiotics were prepared based on the MIC value of each drug. Then, a combination of different dilutions of each drug was prepared in 96-well microtiter plates in a checkerboard formation and the bacterial strain was added. After 20 hrs incubation at 37°C, bacterial growth was measured at the optical density of 620nm. Combinations showing a decrease in the MIC of 90% in comparison to the MICs of the antibiotic and CFTR modulator alone were recorded.. The fractional inhibitory concentration (FIC) index was calculated as $MIC_{90}$ of CFTR modulator in combination/ $MIC_{90}$ CFTR alone + $MIC_{90}$ antibiotic in combination/ $MIC_{90}$ antibiotic alone [5]. If no endpoint MIC value could be determined because of resistance to compound or solubility limits, the next highest MIC value was chosen for the FIC calculation (i.e., if the MIC>32, an MIC=64 was selected for the calculation) [6]. The interaction between drugs was interpreted as synergistic (FIC index <0.5), additive (FIC index $\geq$0.50 and <1.0), indifferent (FIC index $\geq$1.0 and $\leq$4) and antagonistic (FIC index >4) [5].

**Bacteria preparation for acute infection.** An aliquot of *P. aeruginosa* PAO1 reference strain from glycerol stocks was streaked for isolation on TSA and incubated at 37°C O/N. One colony was picked from the plate and used to inoculate 5 ml of TSB and placed in a shaking incubator at 37°C 200 rpm O/N. The O/N bacterial suspension was diluted to 0.1 OD/ml in 20 ml of TSB / flask and grown for 3 hrs at 37°C at 200rpm, to reach the log phase [1]. The bacteria were pelleted by centrifugation (2.700 g, 15 min, 4°C), resuspended in sterile phosphate-buffered saline (PBS) and diluted to give $1x10^6$ colony forming units (CFUs) / mouse in 60 µl.

**Mouse models.** C57BL/6NCrlBR male mice (8-10 weeks of age) were purchased from Charles River (Calco, Italy), shipped in protective, filtered containers, transported in climate-controlled trucks, and allowed to acclimatize for at least two days in the stabulary prior to use. Mice were maintained in sterile ventilated cages in the biosafety level 3 (BSL3) facility at San Raffaele Scientific Institute (Milano, Italia) where 3-5 mice per cage were housed. Mice were fed with standard rodent autoclaved chow (VRFI, Special Diets Services, UK) and autoclaved tap water. Fluorescent lights were cycled 12hrs on, 12hrs off, and ambient temperature (23±1°C) and relative humidity (40-60%) were regulated. For PK experiments, animals were separated into two groups, with IVA levels analyzed in a model of acute *P. aeruginosa* lung infection or in uninfected mice. For infection experiments, mice were anesthetized by an intraperitoneal injection of a solution of Avertin (2,2,2- tribromethanol, 97%) in 0.9% NaCl and administered at a volume of 0.015 ml/g body weight. Mice were placed in supine position. The trachea was directly visualised by ventral midline, exposed and intubated with a sterile, flexible 22-g cannula attached to a 1 ml syringe. An inoculum of 60 µl of planktonic bacterial cells was implanted via the cannula into the lung. After inoculation, all incisions were closed by suture. IVA was administered once, half an hour after the challenge, by intraperitoneal (i.p.) injection at a dose of 3 mg/kg using as vehicle 10% PEG 400, 10%Tween 80, 80% saline. IVA dose used in this model was calculated based on a single adult dose (150 mg) adjusted for mouse weight, assuming that an adult with CF weighs 50 kg [28]. Mice were euthanized with an overdose of carbon dioxide and the diaphragm opened. Blood was collected in 0.5-ml K3-EDTA tubes through cardiac puncture. Time points for sampling were before t = 0.1667, 1, 2, 6, and 24 hrs after administration. Blood samples were directly placed on ice and centrifuged soon after in a precooled centrifuge, and plasma samples were stored at −80°C until IVA plasma concentrations were determined. One sample was taken per mouse, and every time point was sampled in triplicate for uninfected mice (3 mice) and quadruplicate for infected ones (4 mice). Bronchoalveolar lavage (BAL) was performed immediately after blood collection. The concentrations in ELF were determined by taking BAL fluid samples and a concomitant plasma sample, as previously described [37]. In short, murine trachea was exposed by a 1-cm incision on the ventral neck skin for insertion of the cannula. Lungs were instilled two times with 0.9 ml of sterile saline, and the fluid was aspirated immediately. The aspirates recovered from each mouse were pooled and directly placed on ice, then centrifuged in a precooled centrifuge and the supernatant subsequently stored at −80°C. Lungs were excised aseptically, weighed and homogenized in 2 ml of saline using the homogenizer gentleMACS$^{TM}$ Octo Dissociator, and then stored at -80°C.

**Protein binding and concentration determinations in plasma, lung and ELF.** Protein binding was measured in mouse plasma and lung homogenate by Rapid Equilibrium Dialysis (RED). Stock solutions of ivacaftor (IVA) and its main metabolite M1 were prepared in MeOH whereas M6 was prepared in DMSO at a concentration of 1 mg/ml. Working solutions (WSs) were prepared by adding 10µl of each stock solution to 90µl of DMSO to have a concentration of 0.1 mg/ml. The standard Diclofenac, used as control as highly binding standard [7], was prepared in DMSO (1 mM). 5 µl of WSs were diluted in 495 µl plasma or lung homogenate to have a final compound incubation concentrations of 1000 ng/ml and 10 µM for Diclofenac. The concentration of 1000 ng/ml was chosen because similar to IVA $C_{max}$ *in vivo*. Spiked samples (200 µl) were dialyzed in the Rapid Equilibrium Dialysis (RED) inserts (Thermo Scientific) against 350 µL of PBS buffer in the RED plate for 4 hrs at 37°C, under agitation (300 rpm orbital shaker). Samples were tested in duplicate. At the beginning and the end of the incubation, 50 µl of sample in the donor were added to 50 µl of PBS buffer whereas 50µl of the blank matrix were added to 50 µl of dialyzed PBS buffer. Samples were then mixed with 200 µl of acetonitrile containing the internal standard (Lidocaine 10 ng/ml for IVA and metabolites, and Verapamil 10 ng/ml for Diclofenac). Samples were centrifuged at 3500g for 5 minutes. Then, supernatants were transferred into vials for ultra-performance liquid chromatography-tandem mass spectrometry (UPLC-MS/MS) analysis. $T_0$ was used to check potential instability in the matrix. 5 µl supernatant was injected on a Waters Acquity Premiere Ultrahigh-Performance Liquid Chromatography (UPLC) system equipped with a Kinetex C8 (2,6µm particle size, 100A 50*2.1mm Phenomenex) column at 35 °C, running at a flow rate of 300 µl/min and interfaced with an AB Sciex API 3200 mass spectrometer with a TurboIonSpray source (positive mode) and the Analyst (version 1.6.3) software. The chromatographic gradient was in 0.1% formic acid in water or acetonitrile, going from a 2% organic phase at 2 min to 98% at 2.5min. Isocratic condition at 98% was kept up to 4 min and the system was reconditioned from 4.5 to 6 min at 2%B. Mass spectrometer was run in Positive mode with a capillary Temperature 450 C, CUR 30, Gas 1 25 and Gas 2 45, IS was at 5500V. Quantitation was performed in Multiple reaction Monitoring (MRM) using the transitions reported in **Supplementary Table S2**. Protein binding was calculated as follows:

% Fb (fraction bound) = [(area donor – area acceptor) / area donor] *100

Free fraction= 100- %Fb.

Concentrations of IVA and its metabolites in ELF were determined by using the ratio of the urea concentration in BALF to that in plasma [8]. Urea content was measured with a colorimetric Sigma Aldrich MAK006 Kit. Absorbance was measured at 570 nm (A570) on a Tecan Sunrise spectrophotometer. Urea concentrations are normally the same in plasma and ELF because of the equilibrium across the capillary-alveolar membrane. Thus, the apparent ELF volume was estimated by using urea as an endogenous marker of ELF dilution and was calculated as follows: drug concentration in ELF = drug concentration in BALF × urea concentration in plasma/urea concentration in BALF. Reference urea standards ranged from 20 to 100 nmol/ml and were linear over the concentration range.

**Samples preparation.** Lung homogenates were prepared by homogenizing the whole tissue in 2 ml of ammonium acetate buffer 20mM. An intermediate 50 µg/ml stock solution of all the three analytes was prepared from the stock solutions 1 mg/ml. WSs of the three compounds, with concentrations ranging from 5 to 10000 ng/ml, were prepared by sequential dilution of intermediate stock solution in MeOH. Calibration curves in plasma and lung homogenate and BALF were prepared by spiking 5 µl of WSs into 45 µl of plasma, blank lung homogenate or blank BALF. QC samples were analogously prepared at concentrations of 10, 50 and 500 ng/ml in the three matrices. Calibrants, QC samples and samples (50 µl of matrices) were added to a 96 wells plate Phree Phospholipid Removal Solutions (Phenomenex) with 200 ul of acetonitrile containing Lidocaine as internal standard (10 ng/ml) and 0.1 % of formic acid. The plate was shaken for 5 min on a 500g orbital shaker and centrifuged for 15 min at 5°C with a collection plate. The eluates were transferred in a 96-well plate for autosampler and analyzed. Calibration ranges were the following ones: for IVA 0.5-750 ng/ml in plasma and BALF and 4-1000 ng/g in the lung homogenate; for M1 0.5-1000 in plasma, 0.5-500ng/ml in BALF and 3.4-1000 ng/g in the lung homogenate; for M6 0.5-500ng/ml in plasma and BALF, and 3.4-2000ng/g in the lung homogenate. Concentration values reported as unquantifiable were excluded from the evaluation.

**Supplementary Tables**

**Supplementary Table S1. Phenotypic traits and antibiotics susceptibility of clinical *P. aeruginosa* isolates tested in this study.** *P. aeruginosa* early isolates, collected from CF patients at the early stage of chronic colonization, and their pathoadaptive variants, collected after years of persistence, in the advanced stage of chronic colonization (late), were previously characterized for their clonality (all isolates collected from a single patient are clonal), phenotypic traits and susceptibility to antibiotics [3, 9, 10].

| Patient | Isolate / type | Yrs (Δ) | Mucoidy µg uronic acid/ µg protein | Motility | | Protease (ø cm) | Pyocyanin ($OD_{695}$ 26h) | Susceptibility to antibiotics | | | |
|---|---|---|---|---|---|---|---|---|---|---|---|
| | | | | Twitching (ø cm) | Swimming (ø cm) | | | COL | CIP | MER | TOB |
| AA | 2 / early | | - | 1.8 | 5.8 | 1.8 | 0.05 | - | - | - | - |
| | 43 / late | 7.5 | 1.00 | 1.1 | - | 1.0 | 0.02 | - | - | - | - |
| | 44 / late | 7.5 | - | 1.3 | - | 1.1 | 0.05 | - | - | - | - |
| MF | 1 / early | | - | 2.6 | 1.6 | 1.7 | 0.05 | - | - | - | - |
| | 51 / late | 10.1 | - | - | - | - | 0.08 | - | R | R | - |
| KK | 1 / early | | - | 1.4 | 1.2 | - | 0.02 | - | - | - | - |
| | 2 / early | | - | - | - | - | 0.01 | - | - | - | - |
| | 71 / late | 12.6 | - | - | - | - | 0.01 | - | R | R | - |
| | 72 / late | 12.6 | - | - | - | - | 0.01 | - | R | R | - |
| RP | 73 / late | 8.5 | - | - | - | - | 0.05 | | | R | |

Yrs (Δ): years between early and late in clonal sequential isolates; * Indicates swimming and twitching motility zone diameter. R: antibiotic resistance. COL: colistin, CIP: ciprofloxacin, MER: meropenem, TOB: tobramycin

**Supplementary Table S2: Phenotypic traits of clinical *S. aureus* isolates tested in this study.** *S. aureus* early isolates, collected from CF patients at the early stage of chronic colonization, and their pathoadaptive variants, collected after years of persistence, in the advanced stage of chronic colonization (late), were previously characterized for their clonality (all isolates collected from a single patient are clonal) and phenotypic traits [11].

| Patient | Isolate type | Yrs (Δ) | Colony size | CP | β-HL |
|---|---|---|---|---|---|
| 20 | early | | n | +++ | |
| | late | 13 | i | ++ | |
| 22 | early | | n | | |
| | late | 13.5 | i | | |
| 25 | early | | n | + | + |
| | late | 13 | n | | |

Yrs (Δ): years between early and late in clonal sequential isolates; n: normally-sized colonies, i: intermediate-sized colonies; CP: capsule polysaccharide production, β-HL: presence of β-haemolysin activity.

**Supplementary Table S3. Multiple Reaction Monitoring (MRM) transitions and conditions**

| Analyte | Parent Ion MH+ (Da) | Product Ion (Da) | Dwell Time (msec) | Declustering Potential (Volts) | Collision Energy(volts) |
|---|---|---|---|---|---|
| IVA_1* | 393.2 | 172.2 | 100 | 38 | 38 |
| IVA_2 | 393.2 | 319.2 | 100 | 43 | 41 |
| M1_1* | 409.1 | 172.3 | 100 | 44 | 48 |
| M1_2 | 409.1 | 353.0 | 100 | 44 | 23 |
| M6_1* | 423.1 | 172.3 | 100 | 45 | 41 |
| M6_2 | 423.1 | 367.2 | 100 | 45 | 23.5 |
| Lidocaine 1* | 235.3 | 86.1 | 100 | 49 | 26 |
| Lidocaine 2 | 235.3 | 58.1 | 100 | 44 | 52 |

*MRM transitions used for quantitation

**Supplementary Table S4. Minimum inhibitory concentrations ($MIC_{90}$) of ivacaftor (IVA), lumacaftor (LUM), tezacaftor (TEZ), elexacaftor (ELX) and the triple combination elexacaftor/tezacaftor/ivacaftor (ELX/TEZ/IVA) against *P. aeruginosa* isolates collected from CF patients at different stages of colonization (early and late).** *P. aeruginosa* isolates were grown for 20 hours in the presence of serial dilutions of IVA, LUM, TEZ, ELX and the triple combination ELX/TEZ/IVA. The concentrations tested ranged from 0.25 µg/ml to 32 µg/ml for IVA, LUM, TEZ and ELX, and from ELX 0.5 µg/ml / TEZ 0.25 µg/ml / IVA 0.375 µg/ml to ELX 64 µg/ml / TEZ 32 µg/ml / IVA 48 µg/ml for the triple combination. The $MIC_{90}$ was defined as the lowest compound concentration showing a reduction in the optical density at 620 nm of approximately 90% in comparison to the optical density of the bacteria grown with the vehicle after 20 hrs. Each experiment was performed at least two independent times (two technical replicates).

| | | ***P. aeruginosa* isolates** | | | | | | | | | | |
|---|---|---|---|---|---|---|---|---|---|---|---|---|
| **Isolate name** | | **PAO1** | **AA2** | **AA43** | **AA44** | **MF1** | **MF51** | **KK1** | **KK2** | **KK71** | **KK72** | **RP73** |
| Isolate type | | ref | early | late | late | early | late | early | early | late | late | late |
| **$MIC_{90}$** (µg/ml) | **IVA** | >32 | >32 | >32 | >32 | >32 | >32 | >32 | >32 | >32 | >32 | >32 |
| | **LUM** | >32 | >32 | >32 | >32 | >32 | >32 | >32 | >32 | >32 | >32 | >32 |
| | **TEZ** | >32 | >32 | >32 | >32 | >32 | >32 | >32 | >32 | >32 | >32 | >32 |
| | **ELX** | >32 | >32 | >32 | >32 | >32 | >32 | >32 | >32 | >32 | >32 | >32 |
| | **ELX / TEZ / IVA** | >64 / >32 / >48 | >64 / >32 / >48 | >64 / >32 / >48 | >64 / >32 / >48 | >64 / >32 / >48 | >64 / >32 / >48 | >64 / >32 / >48 | >64 / >32 / >48 | >64 / >32 / >48 | >64 / >32 / >48 | >64 / >32 / >48 |

ref, reference strain; early, isolate collected at the early stage of chronic colonization; late, isolate collected after years of persistence, in the advanced stage of chronic colonization.

**Supplementary Table S5. Pharmacokinetic parameter estimates for M1 in plasma, epithelial lining fluid (ELF) and lungs of infected and non-infected mice.** C57BL/6NCrlBR male mice (8 to 10 weeks of age) were infected with 1x10$^6$ colony forming units of *P. aeruginosa* PAO1 by intratracheal administration. A non-infected control group was also tested in parallel. After 30 min from the infection mice were treated with 3 mg/kg ivacaftor (IVA) in 10% PEG 400, 10%Tween 80, 80% saline by intraperitoneal administration. Mice were sacrificed at 10 min, 1, 2, 6 and 24 hrs after IVA administration. Blood was collected and processed to obtain plasma. Bronchoalveolar lavage fluid (BALF) was collected, centrifuged and the supernatant used to quantify IVA concentration. Lungs were excised, homogenized, centrifuged and the supernatants used to quantify IVA concentration. Plasma, lung homogenate and BALF (50µl) were added to a Phree Phospholipid Removal plate (Phenomenex) with acetonitrile and 0.1% formic acid in order to eliminate phospholipids, decreasing the matrix effect. Eluates were analyzed by UPLC-MS/MS method with a linear gradient in MRM positive mode. Calibration ranges were the following ones: M1 0.5-1000 in plasma, 0.5-500ng/ml in BALF and 3.4-1000 ng/g in the lung homogenate. Data are the geometric mean of values from 3-4 mice. Concentrations of M1 in ELF were determined using the ratio of the urea concentration in plasma to that in BALF. Concentration in ELF = drug concentration in BALF × urea in plasma / urea in BALF. The data are the geometric means of values from 3-4 mice.

| Sample | Mice group | Pharmacokinetic parameters | | | | | | | |
|---|---|---|---|---|---|---|---|---|---|
| | | $T_{max}$ (hrs) | $C_{max}$ (ng/ml) | $T_{last}$ (hrs) | $C_{last}$ (hrs) | $AUC_{last}$ (hrs ng/ml) | $AUC_{INF}$ (hrs ng/ml) | $T_{1/2}$ (hrs) | MRT (hrs) |
| plasma | non-infected | 1 | 111.6 | 24 | 2.0 | 887.3 | 898.7 | 4,0 | 4.9 |
| | infected | 1 | 82.6 | 24 | 1.6 | 823.3 | 832.5 | 4,0 | 5.0 |
| ELF | non-infected | 6 | 10.8 | 6 | 10.8 | 55.1 | nd | nd | 3.3 |
| | infected | 1 | 11.6 | 6 | 9.9 | 59.9 | 406.6 | 24.3 | 3.1 |
| lung | non-infected | 2 | 7.0 | 6 | 6.4 | 36.3 | nd | nd | 3.2 |
| | infected | 6 | 18.4 | 6 | 1.7 | 239.8 | nd | nd | 6.7 |

$T_{max}$, time of maximum concentration; $C_{max}$, maximum concentration; $T_{last}$, time of last quantifiable concentration; $C_{last}$, last quantifiable concentration; $AUC_{last}$, AUC to the last quantifiable concentration level; $AUC_{INF}$, AUC to infinity; $T_{1/2}$, half-life; MRT, mean residence time; nd, not determined.

**Supplementary Table S6. Penetration of M1 in lung and epithelial lining fluid (ELF).** Plasma and lung concentrations were multiplied for the free fraction from the protein binding experiment (0.11% for plasma and 2.83% for lung; **Table 4**). ELF free fraction was calculated considering ELF protein binding equal to protein binding in the bronchoalveolar lavage fluid. Data are the geometric mean of values from 3-4 mice.

| Mice group | Values by sample type | | | | | | lung/plasma | | ELF/plasma | |
|---|---|---|---|---|---|---|---|---|---|---|
| | plasma | | lung | | ELF | | | | | |
| | $fC_{max}$ (ng/ml) | fAUC (hrs ng/ml) | $fC_{max}$ (ng/ml) | fAUC (hrs ng/ml) | $C_{max}$ (ng/ml) | AUC (hrs ng/ml) | $C_{max}$# | AUC# | $C_{max}$# | AUC# |
| non-infected | 0.12 | 0.97 | 0.20 | 0.93 | 10.8 | 49.0 | 1.67 | 0.96 | 90.0 | 50.5 |
| infected | 0.09 | 1.10 | 0.95 | 1.57 | 11.6 | 52.0 | 10.6 | 1.43 | 129 | 47.3 |

### Values for lung/plasma (ratio of lung free maximum concentration or exposure to plasma free maximum concentration or exposure) and ELF/plasma (ratio of ELF maximum concentration or exposure to plasma free maximum concentration or exposure); $fC_{max}$ and fAUC refer to unbound M1 in plasma and lung.

**Supplementary Figure**

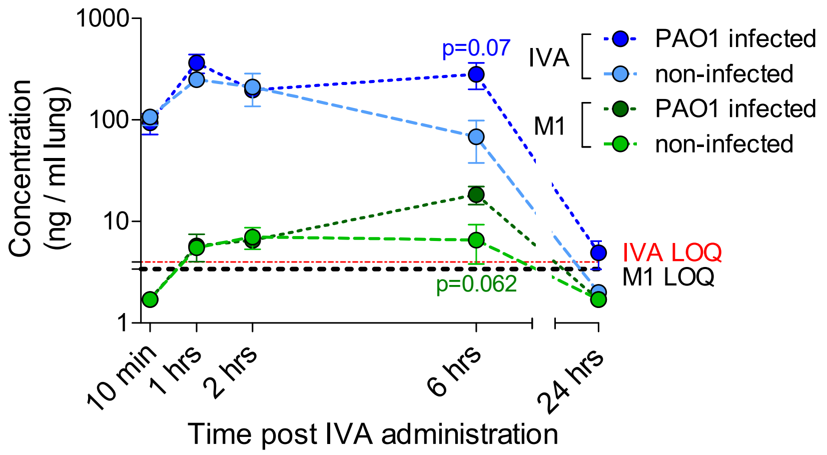


**Supplementary Figure S1. Ivacaftor (IVA) and M1 concentrations in murine lung.** C57BL/6NCrlBR male mice (8 to 10 weeks of age) were infected with $1x10^6$ colony forming units of *P. aeruginosa* PAO1 by intratracheal administration. An uninfected control group was also tested in parallel. After 30min from the infection mice were treated with 3 mg/kg IVA in 10% PEG 400, 10%Tween 80, 80% saline by intraperitoneal administration. Mice were sacrificed at 10min, 1, 2, 6 and 24 hrs after IVA administration. After blood and bronchoalveolar lavage fluid collection, lungs were excised, homogenized and the supernatants used to quantify IVA and M1 by high-performance liquid chromatography–tandem mass spectrometry. Data, derived from 3-4 mice, are represented as mean values ± standard errors of the means (SEMs). Limits of quantification (LOQ) for IVA and M1 are indicated. A value corresponding to LOQ/2 was assigned to undetectable samples at specific time-points (10 min and 24 hrs). Statistical significance was calculated by Mann-Whitney test comparing infected and uninfected mice at each time-point.
